# Recombinant binder of sperm protein1 (rec-BSP1) as a potent fertility factor mediates its effect on spermatozoa and production of blastocysts in buffalo

**DOI:** 10.1101/2021.04.12.437362

**Authors:** Sudam Bag, Shama Ansari, Alka Turuk, Nimai Charan Mahanandia, Sikander Saini, Satya Kumar Biswal, Sandeep Kumar Singh, Praveen Malik, Satish Kumar, Dhruba Malakar

## Abstract

**Objective:** To understand the effect of recombinant BSP1 (rec-BSP1) on *in vitro* capacitation of sperm and fertilization study

**Method(s):** Articles were screened for reports including rec-BSP1, Capacitation, *in vitro* fertilization

**Intervention:** None

**Main Outcome Measure(s):** Reproductive outcomes, effect on gametes and embryos

**Result(s):** Here we report an optimization of condition for rec-BSP1 production which was used for *in vitro* capacitation and enhancement of buffalo embryo production. The sequence of the protein was used for multiple sequence alignment which has 99% similarity with PDC 109 protein. The expression of rec-BSP1 was carried out successfully with 1 mM IPTG at 16^0^ C for 22 hrs and purified it in soluble form. The structure of rec-BSP1 was generated using 3D modelling and analysed its mode of binding with heparin and PC by molecular docking and the structural stability of rec-BSP1-PC and rec-BSP1-heparin complexes by using molecular dynamic (MD) simulation. The effect of rec-BSP1 was observed on *in vitro* capacitation of spermatozoa and buffalo blastocyst production. It was found that the rec-BSP1 enhanced the sperm motility at a concentration of 50 μg/ml for 1 h of incubation without having any detrimental effect on the sperm morphology and a significant increase in blastocyst production at concentration of 50 μg/ml rec-BSP1. Hence this finding represents a new insight and advance the prospective approach to develop a potential fertility factor in reproduction.

**Conclusion(s):** The purified rec-BSP1 may enhance on male fertility and mediated its effect on *in vitro* blastocyst production in buffalo.

---

Binding of sperm proteins (BSPs) (1) one of the major proteins of seminal plasma in male that contributes around 70% of total seminal plasma proteins (2). The BSPs have a unique structure and are involved diverse functions such as maintenance of survivability, motility, fertility of spermatozoa (3–10), capacitation (11), fertilization (12) and protective action against stressful conditions (13). It is believed that it plays a significant role in selection of high-quality sperms for successful fertilization. In general, all isoforms of BSP protein are having two fibronectins type II (FN2) domains separated by polypeptide linker. Each FN2 domain of BSPs is conserved among the family (14) which binds to different ligands like heparin, PC, gelatine, HDL, glycosaminoglycan (15–18). The binding induces conformational changes in the BSPs, which in turn induce cholesterol efflux, sperm capacitation and trigger acrosome reaction (19). The association of different ligands with BSPs has widely been studied to investigate the effect of fertility in males (20). However, the ligand-binding mechanism for rec-BSP1 and the function of rec-BSP1–ligand complexes are poorly explored. The native purification of PDC-109 protein from buffalo was done by conventional method which is antigenically similar to BSP protein in buffalo (21, 22). The availability of pure BSPs can help in the studies related to improvement of fertilization and cryopreservation of spermatozoa of buffalo for faster multiplication of superior germplasm. The paucity of tissue sample is a main hurdle in the purification of BSPs in its native form. The recombinant expression of BSPs is an appropriate alternative to the native pure proteins provided the recombinant proteins are properly folded. There are no available reports on the use of buffalo rec-BSP1 in *in vitro* capacitation and embryo production. Therefore, the present study was envisaged to understand the mode of binding of buffalo rec-BSP1 on heparin and PC which play a significant role in capacitation of spermatozoa and effect on *in vitro* embryo production (23). The several computational approaches such as molecular modelling, docking, MD simulation, and binding free energy estimation were taken into consideration to understand the structural integrity of rec-BSP1 and mode of binding with these ligands. Keeping in view of lower rate of fertility in buffaloes, the present investigation has been conducted to develop a method for production of recombinant BSP in E. coli in soluble form and establishing its role in modulation of sperm morphology and function and evaluated its effect on *in vitro* buffalo embryo production.

## OBJECTIVE

The aim of this review is currently known about the effect of rec-BSP1 on capacitation of sperm, *in vitro* production of embryos in buffalo

## METHODS

### Retrieval Sequence of BSP1

The nucleotide sequence of binder of sperm protein1 (BSP1) (*<u>Bubalus bubalis</u>*) (Accession Number: KR703587.1) was retrieved from NCBI database. The nucleotide sequence was optimized and translated into protein sequence by using EMBOSS Transeq tool. This protein sequence was designated as recombinant binder of sperm protein (rec-BSP). For sequence analysis, the protein sequence of rec-BSP having similar sequence of other species such as human, mouse, pig, canine, horse and bovine were taken from UniProt database. Multiple sequence alignment between rec-BSP1 with selected sequences was performed by ClustalW2 program. Further, phylogenetic tree was constructed using Mega X software. In Mega X software, the sequences were aligned by muscle program using UPGMA method. The phylogeny tree was constructed by using maximum likelihood method (ML) based on Jones-Taylor-Thornton (JTT) model with bootstrap value of 1000. Five numbers of discrete gamma distribution categories were taken into account for evolutionary rates using all sites. The tree was obtained by heuristic search using Nearest-Neighbor-Interchange (NNI) method; however, the initial tree was obtained by Neighbor Joining method in MEGA X program.

### Homology Modelling of rec-BSP1

The homology model of rec-BSP1 was built with the Modeller version 9.23 program. For identify suitable template for computational modeling of rec-BSP1, BLASTp tool against PDB (Protein Data Bank) database was used for identifying a suitable template of rec-BSP1. Modeller 9.23 facilitated for development of 50 raw models of which the models with the lowest discrete optimized protein energy (DOPE) score was selected. Further, the three-dimensional (3-D) structure was refined by ModRefiner algorithm (Dong Xu and Yang Zhang, 2011). The final model was validated by PROCHECK and model quality assessment and energy profile characterization was done by ERRAT and ProSA. For correctness and accuracy of 3D modeled of rec-BSP1 protein, 100 ns molecular dynamics simulation was performed using AMBER99SB-ILDN force field in Gromacs version 5.0.7 software.

### Selection of ligands

The two-dimensional (2-D) structures of heparin and phosphatidylcholine (PC) were retrieved from PubChem and ChEBI database. The 2-D structures were converted to a three-dimensional (3D) structures using OpenBabel program (O’Boyle NM, Banck M., 2011). The generated structures were then subjected to energy minimization by using the amber ff14sb force field in the UCSF Chimera tool. Further, these molecules were prepared for molecular docking with rec-BSP1.

### Molecular Docking

The modeled structure of rec-BSP1 and two molecules (Heparin & PC) were used as target and ligands respectively. The molecular docking was performed by using AutoDock 4.2.6 tool. The polar hydrogen and Kollman charges were applied to the structure of rec-BSP1. Geister partial charges were applied for both ligands. The 3D grid box with dimensions X = 70A^0^, Y = 70A^0^, and Z = 70A^0^ with grid-point center X = 40.044A^0^, Y = 18.668A^0^, Z = 25.432A^0^ and the grid spacing of 0.375A^0^ were made to cover the binding cavity of rec-BSP1. The Lamarckian genetic algorithm (LGA) was employed to perform molecular docking in AutoDock. During the docking process, a maximum of 50 conformers were considered for each ligand which was set to terminate after 2500000 energy evaluations. The best docking conformation of the rec-BSP1 and ligands was visualized by Pymol.

### MD simulation of rec-BSP1 and selected ligands

The best binding mode of rec-BSP1 and ligands complexes were subjected to molecular dynamics simulation by GROMACS version 5.0.7 package. MD simulations were performed using GROMOS96 53a6 force field method (Oostenbrink et al., 2004). The heparin and PC topologies files were generated using PRODRAG web server. The protein-ligands complexes were solvated with TIP3P water model and a cubic box setting was done with a distance of 10 A^0^ between the edge of the box and protein surface. Further neutralization of the protein-ligands complex by Na^+^/Cl^−^ ions with 0.15M ionic strength and energy minimization was done using steepest descent algorithm. After energy minimization, both complexes were subjected to equilibration using NVT (constant number of atoms, volume, and temperature) and NPT (constant number of atoms, pressure, and temperature) ensemble 300K for 1 ns. Finally, MD simulation of both complexes were run for 25 ns at a constant temperature (V-rescale method) and pressure (Parrinello-Rahman method) for trajectory analysis. Linear Constraint Solver (LINCS) algorithm was applied to constrain the covalent bonds and partial mesh Ewald method was used for calculating the electrostatic interactions. The cut-off radii for the coulomb and van der Waals interactions were fixed at 10.0 A^0^ and 14.0 A^0^. The MD trajectories snapshots saved at the interval of 100 ps during the simulation period. To analyze the MD trajectories, GROMACS built-in modules utilities like root-mean-square deviation (RMSD), Radius of gyration (RG), Residue-based Root Mean Square Fluctuations (RMSF), Solvent accessible surface area (SASA) and inter-molecular hydrogen bond (H-Bond) were used. The 2D graphs were plotted using Xmgrace.

### Conformation of PCR products

The optimized nucleotide sequence was amplified by one primer set designed in primer-blast software using Binder of sperm 1 (BSP1) mRNA (*<u>Bubalus bubalis</u>*), complete CDS (KR703587) with standardized annealing temperature of 52^0^C and confirmed their size on 1% agarose gel. (F-5’AGCTGGATCCATGGCACTGCAGTTGGGGCTC 3’R-5’CTAGCTCGAGCTAATGGTGGTGGTGATGATGGCAATACTTCCAAGCTCTG TCCTT3’).

### Sub-cloned in Expression vector and screening of recombinant plasmids

In order to express BSP1, the expression vectors and recombinant plasmids were digested with *NdeI* and *XhoI* restriction enzymes. The purified inserts and vector were ligated in 3:1 molar ratio (24). The recombinant plasmids were transformed in E. Coli cell BL21 (DE3) for expression. Screening of the recombinant colonies was carried out by colony PCR and digested the plasmid DNA with *NdeI* and *XhoI* restriction enzymes for insert release (25).

### Purification and confirmation of rec-BSP1

The expression conditions of the protein like IPTG concentration, temperature and incubation period were optimized for production of desired amount of rec-BSP1 in soluble form (26). The recombinant protein was expressed as 6x His tag fused to N-terminal of proteins hence it was purified by Ni-NTA agarose column. The protein was eluted from beads using pH 4.5 buffers and purity of proteins checked on SDS-PAGE (27). Further the protein was confirmed by Western Blot and quantified by Braford assay.

### *In Vitro* Capacitation study

Semen samples were collected from Animal Breeding Research Center (ABRC), National Dairy Research Institute and 0.1ml of semen having concentration of 2-4 million spermatozoa/ml was transferred to 9 ml of working Brackett and Oliphant (BO) (28) medium and centrifuged at 1200 rpm for 7 min. 2-3 washing was done using same BO medium. The resulting pellet was dissolved in BO medium and distributed in two groups (treatment & control) which supplemented with concentration of 50 μg/ml rec-BSP1 and heparin at concentration of 20 μg/ml for incubation in a humidified CO_2_ incubator (5% CO_2_, 95% RH) at 38.5^0^C for 1-4 hr. The progressive motility of spermatozoa assessed by CASA and staining (Eosin, nigrosine & trypan blue) was done for morphology of spermatozoa. Similarly, Hoechst staining was performed to assess the acrosome status of spermatozoa (29) and observed it under oil immersion lens (100X) using light microscope to evaluate the proportion of acrosome reactions.

### *In Vitro* production of embryos in buffalo

The effect of rec-BSP1 on *in Vitro Fertilization* study was done by treatment of different concentration (20, 50 & 100 μg/ml) of the recombinant protein and heparin at concentration of 20 μg/ml in fertilization BO medium. The processed spermatozoa and oocytes were co incubated for cleavage, blastocyst and hatched blastocysts formation. Ovaries were collected from slaughter house (Gazipur, Delhi) and processed for maturation in presence of exogenous hormonal supplement. COCs were cultured in group of 15-20 nos in 100 μl droplets of the IVM medium (TCM-199+10 % FBS+5μg/ml pFSH +1μg/ml estradiol-17β+0.81mM sodium pyruvate+50 μg/ml gentamycin sulfate) and overlaid with sterile mineral oil in Petri dishes for 24 hr in a humidified CO_2_ incubator (5% CO_2_, 95% RH) at 38.5^0^C. The frozen semen was processed for capacitation of spermatozoa (2-4 million spermatozoa/ml) in fertilization BO medium supplemented with 20, 50, 100 μg/ml rec-BSP1 and heparin incubated with matured oocytes for 16-18 hrs in a humidified CO_2_ incubator (5% CO_2_, 95% RH) at 38.5^0^C for *in Vitro Fertilization*. The presumptive zygotes were cultured on original beds of granulosa cells using media mCR2aa supplemented with different growth factors such as IGF-1, EGF, SCF to enhance the development of embryos into late blastocyst and hatching stage (30). Similarly, cysteamine, melatonin, ascorbic acid, mercapto ethanol and glutathione etc. were added to protect embryos from oxidative stress and cultured up to 8 days’ post insemination in a humidified CO_2_ incubator (5% CO_2_, 95% RH) at 38.5^0^C.

## RESULTS

### Analysis of BSP Sequence

The nucleotide sequence of mRNA of BSP (Bubalus bubalis) was retrieved from NCBI database. The nucleotide sequence was optimized by using Codon-Opt software and translated into protein sequences by EMBOSS Transeq tool. The similar BSP protein sequences of different species (human, mouse, pig, canine, horse and bovine) were taken from UniProt database for multiple sequence alignment by using ClustalW2 program (Supplementary fig. S1). The sequences were aligned by Muscle program using UPGMA method. MegaX software was used to reconstruct the phylogenetic tree by maximum likelihood (ML) method and LG model with 1000 bootstrap replicates (Supplementary fig. S2). It was shown in the phylogenetic tree, the recombinant protein having 99% similarity with bovine seminal plasma protein PDC 109 (OS: Bos Taurus).

### Three Dimensional Modeling of rec-BSP1 and structure Validation

Modeller v9.23 software was used to predict the three dimensional (3D) structure of rec-BSP1 sequence. Identification of template through BLASTp search reveals that, the structure of seminal plasma protein PDC-109 (*Bos taurus*) with PDB ID: 1H8P, Chain A with 81% query coverage, 86.73% sequence identity and structure of KDA type IV collagenase (*Homo sapiens*) with PDB ID: 1EAK, Chain A with 60 % query coverage, 34.69% sequence identity were selected as templates for three dimensional (3D) modelling of the rec-BSP1 sequence. Modeller 9.23 employed to develop 50 raw models, the model with the lowest DOPE score and RMSD value was selected for further refinement. Galaxy Loop refinement protocol was applied to refined the loops. The refined model of rec-BSP1 was validated through various protein structure validation server. Ramachandran plot analysis for structural validation was confirmed that the model has 111 (93.3%) amino acid residues in the favored regions, 6 (5%) amino acid residues in additional allowed regions, 2 (1.7 %) in generously allowed regions and 0 (0%) amino acid residue in the disallowed region. The Z-score (−3.91) for rec-BSP1 was calculated by Pro-SA web which is well within the range of experimental structure (Fig. 1 A, B). The validation of secondary structure of rec-BSP1 carried out by STRIDE web server (Fig. 1 C, D). For correctness and accuracy of 3D modeled of the rec-BSP1 protein, 100 ns MD simulation was performed using AMBER99SB-ILDN force field in Gromacs version 5.0.7 software (Fig.1 E, F, G). The rec-BSP1, the RMSD value was evaluated in order to check the stability of the complexes and it was shown that after a period of 100 ns indicating stable conformation of the modelled protein.

**Figure1.**
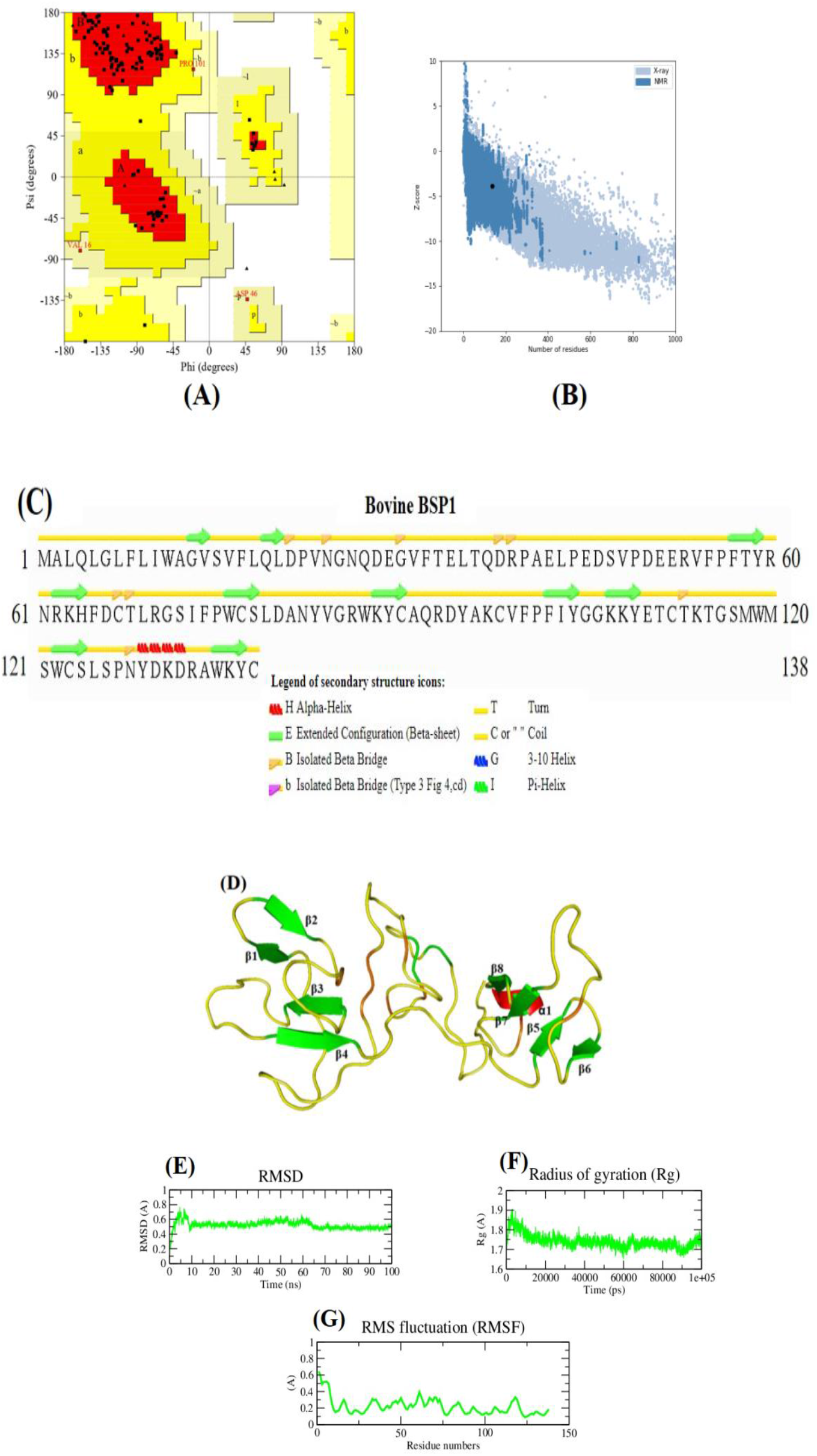
3D Structural model of rec-BSP1. (A) Overall quality of rec-BSP1 structure validation by using procheck for ramachandran plot. (B) rec-BSP1 structure validation using Pro-SA web server. (C) Secondary structure prediction of rec-BSP1. (D) 3D structure of rec-BSP1, the image was generated using PyMOL, where green color represent the Beta-Sheet and red color represent the Coil. (E) graphical representation of RMSD analysis of 100 ns simulation. (F) Radius gyration of 100 ns simulation. (G) RMSF analysis of 138 amino acid residues of rec-BSP1.

**Figure 2.**
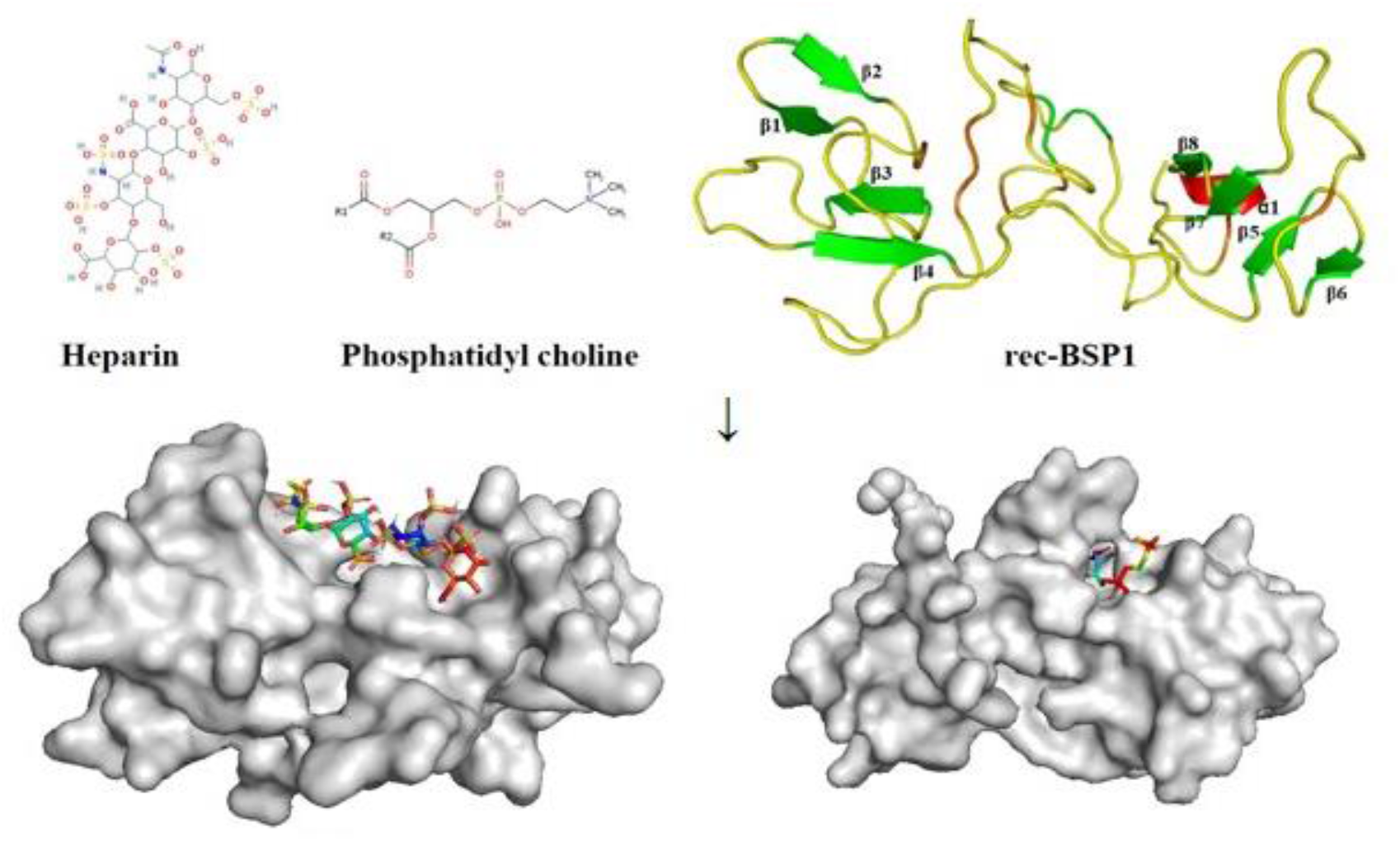
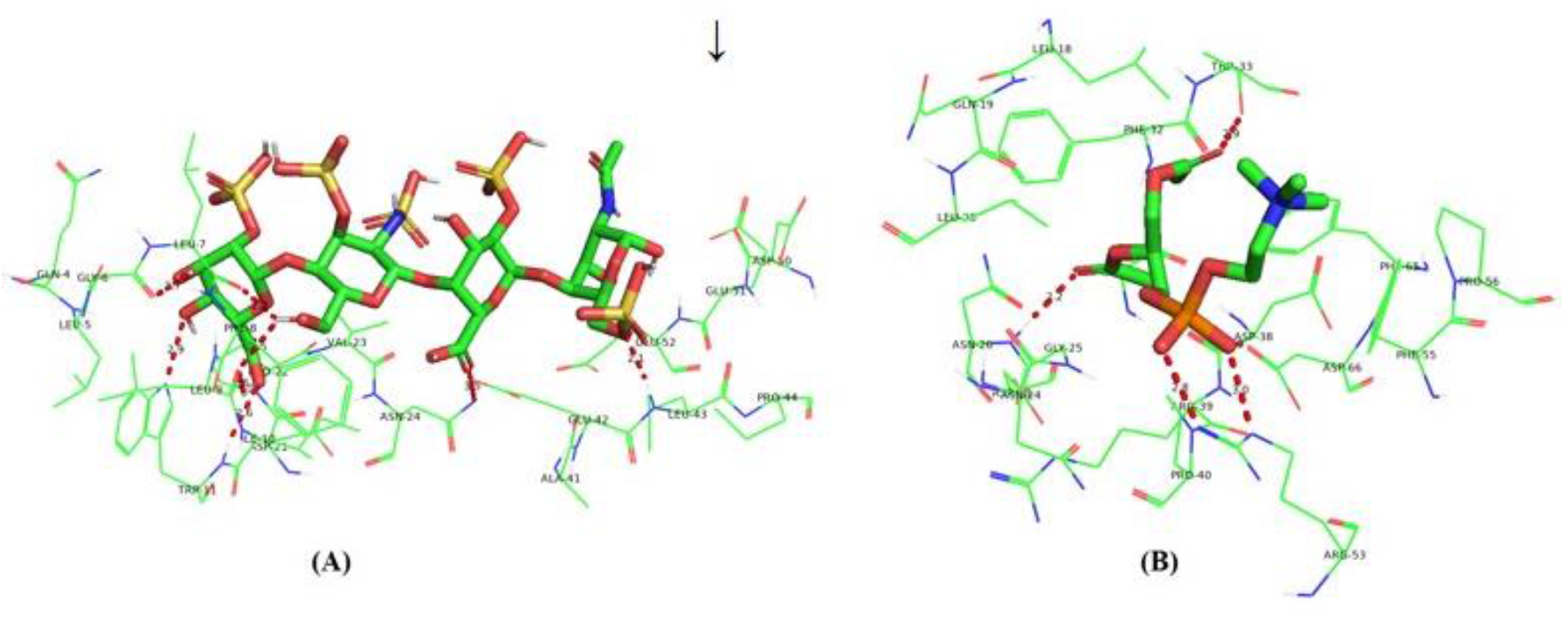
Docking pose of rec-BSP1-ligand complex. The structural representation the binding site of heparin and PC docked complex with interaction represented in red dotted lines. (A) Heparin docked complex. (B) PC docked complex.

### Molecular Docking Analysis

Two ligands were docked to rec-BSP1 using AutoDock 4.2.6 tool. The docking score was determined based on their binding energy, ligand efficiency (31) score and number of hydrogen bonds formed. Summary of the highest binding energy, ligand efficiency and number of hydrogen bonds formed of the ligands is presented (Supplementary Table. S3). The binding site of rec-BSP1 was predicted by metapocket 2.0 web-server (Bingding *et al.,* 2009). The binding site of rec-BSP1 protein residues are ILE10, PHE17, LEU18, GLN19, LEU20, ASP21, VAL23, ASN24, GLY25, ASN26, GLN27, ASP28, GLU29, GLY30, VAL31, PHE32, THR33, GLU34, ASP38, ARG39, PRO40, GLU52, ARG53, VAL54, PHE55, PRO56, PHE65, and ASP66. These residues were selected during molecular docking preparation. The rec-BSP1-heparin complex has binding affinity −7.4 kcal/mol, with four conventional H-bonds whereas rec-BSP1-PC complex has binding affinity −7.4 kcal/mol, with four conventional H-bonds. The rec-BSP1-heparin and rec-BSP1-PC complexes were visualized through PyMol.

### MD Simulation of Heparin and Phosphatidyl Choline docked Complexes

The MD simulation was performed for the heparin and PC complex systems to understand the structural and dynamic stability of the rec-BSP1 when bind with the ligands. The intrinsic dynamical stability of the heparin and PC complex was studied by the root-mean-square deviation (RMSD), radius of gyration (R_g_), solvent accessible surface area (SASA) and root-mean-square fluctuation (RMSF) of the C_α_ atom as a function of simulation time. RMSD measures the difference between the back bone of a protein from its initial structure to its final positions. The stability of protein-ligand complex relative to its preliminary state can be assessed by plotting the deviations produced over the course of simulation. The smaller deviation shows the more stability of the protein-ligand complexes. MD simulation was done to calculate the RMSD value in order to check stability of the complexes and it was shown that average deviation of heparin and PC complexes are approximately 0.43 and 0.89 nm during the course of 25 ns MD simulation. It is indicating that heparin is more stable than PC complex (fig. 3 A). Similarly, RMSF was carried out to understand flexibility and superimposed graphical representation of each residue. The RMSF value was calculated the magnitude of fluctuation of each residue and it was shown that the first complex with heparin having fluctuation range of approximately 0.4–0.7 nm in first 30 residues after that it showed average fluctuation of 0.4 nm. Compared to heparin complex, the plot for PC Complex contain more noises and larger RMSF values, which indicates relatively a weak binding between the target and the substrates. The binding site residues predicted by Metapocket server which were all found to have less fluctuation for the rec-BSP1-heparin complex compared to rec-BSP1-PC complex indicating intactness and rigidity of the binding cavity (fig. 3 B). Radius of gyration value was calculated the compactness and structural changes of complex and observed that the average Rg value of 1.93 nm over the simulation period of 25 ns in the rec-BSP1-ligands complexes. The difference between the rec-BSP1-heparin complex relatively more stable than rec-BSP1-PC complex after a period of 7 ns (fig. 3 C). Analysis of SASA was done for the rec-BSP1 with heparin and PC complexes to understand their interaction and solvent accessibility. The average sasa value for heparin complex was 112, whereas PC complex 107 nm^2^ over a period of 25 ns MD simulation. It was shown that rec-BSP1-heparin complex has less fluctuation than rec-BSP1-PC complex (fig. 3 D). The analysis of hydrogen bond (34) between rec-BSP1 and the ligands were calculated for overall stability of protein-ligand complexes using gmx h-bond over 25 ns simulation period. The rec-BSP1-heparin complex having higher numbers of H-bond (~2.229) compared to rec-BSP1-PC complex (~0.638) (fig. 4 A). It indicates that rec-BSP1-heparin complex was more stable interacting complex. The MD trajectories result files were used for Principal component analysis (PCA) for determination of conformational changes w.r.t rec-BSP1 function. The eigenvalues were calculated the movement of atoms meant for conformational changes which were obtained by diagonalzing the covariance matrix of the Cα atomic fluctuations against the equivalent Eigenvectors indices. It was shown that the projection of trajectory along PC1 and PC2 and rec-BSP1-heparin has a higher value of trace of covariance matrix than PC indicating higher flexibility and increased collective motion. The rec-BSP1-heparin complex showed the least conformational changes due to decrease in collective motions than PC system. The PCA and RMSF investigations evoke similar change about structural fluctuations of the rec-BSP1-ligands complexes (fig. 4 B, C). DSSP algorithm was applied to check the secondary structure protein. It was observed that there was significant structural changes i.e. increase in helical content and absence of *β*-sheet in rec-BSP1 (fig. 4 D, E).

**Figure 3.**
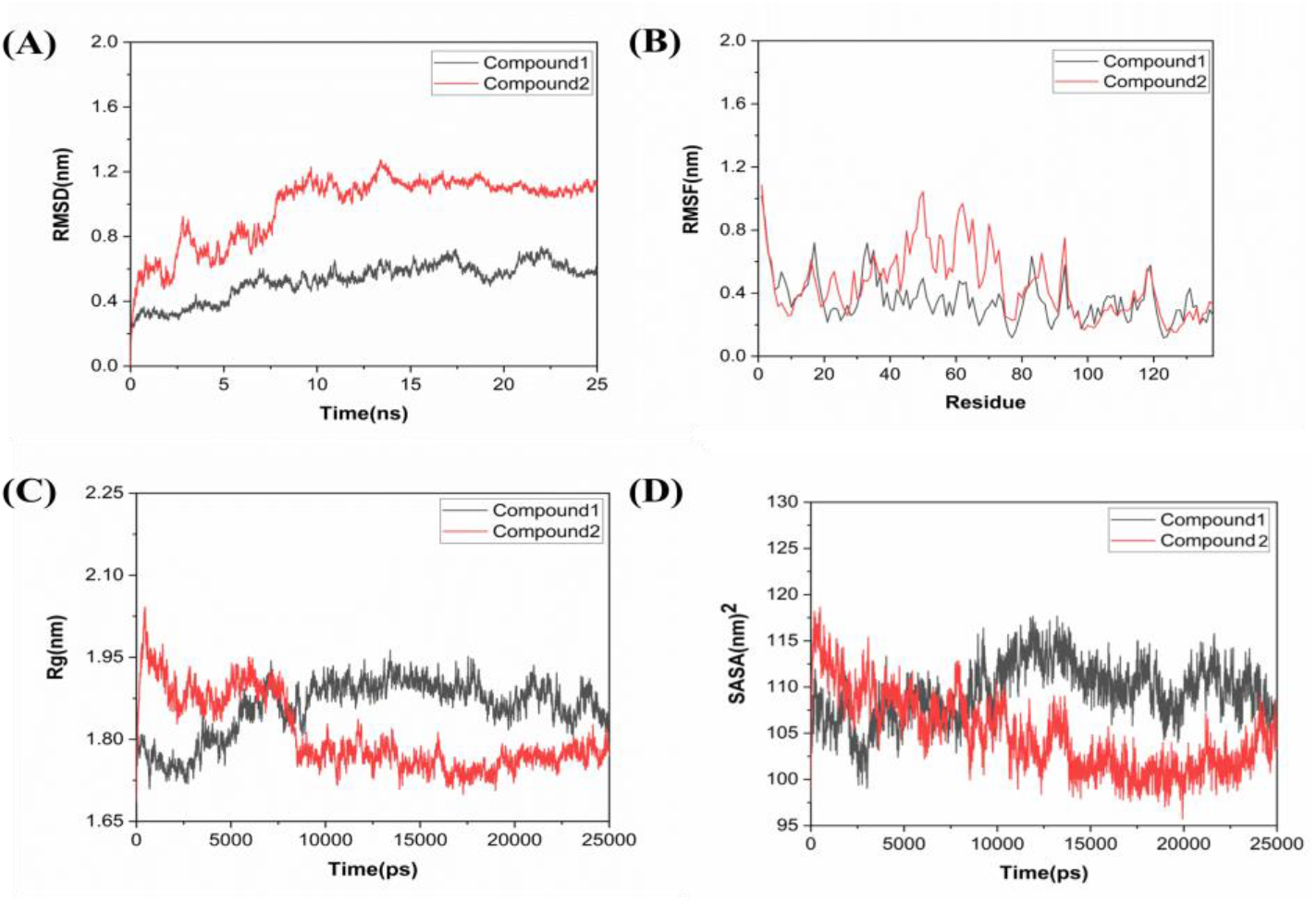
Dynamics stability of rec-BSP1 with heparin and PC complexes during 25 ns MD simulation. (A) RMSD analysis of ligand Compound1 (heparin, black), and Compound2 (PC, red) protein backbone conformation. (B) RMSF analysis of 138 amino-acid residues is represented for Compound1 (heparin, black) and Compound2 (PC, red) protein complexes. (C) Radius gyration of Compound1 (heparin, black) and Compound2 (PC, red). (D) Solvent accessible surface area (SASA) of Compound1 (heparin, black) and Compound2 (PC, red).

**Figure4.**
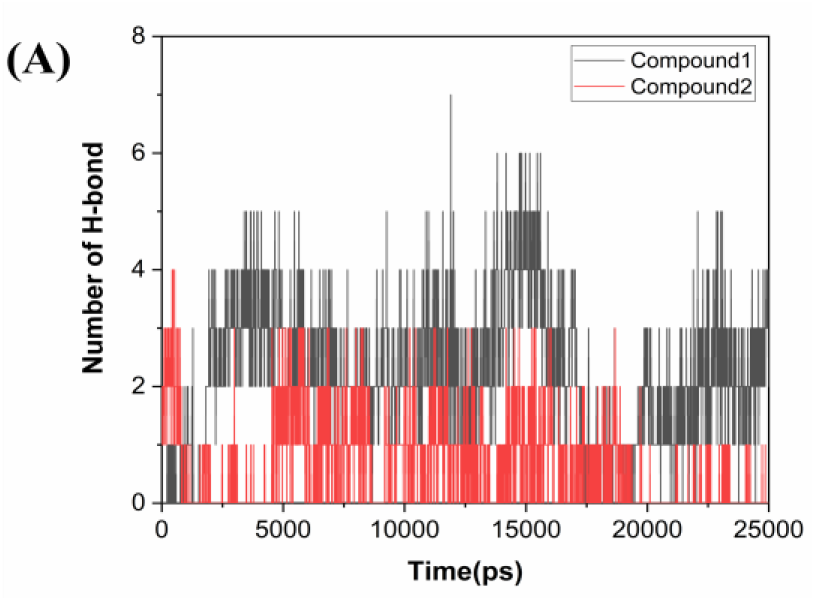
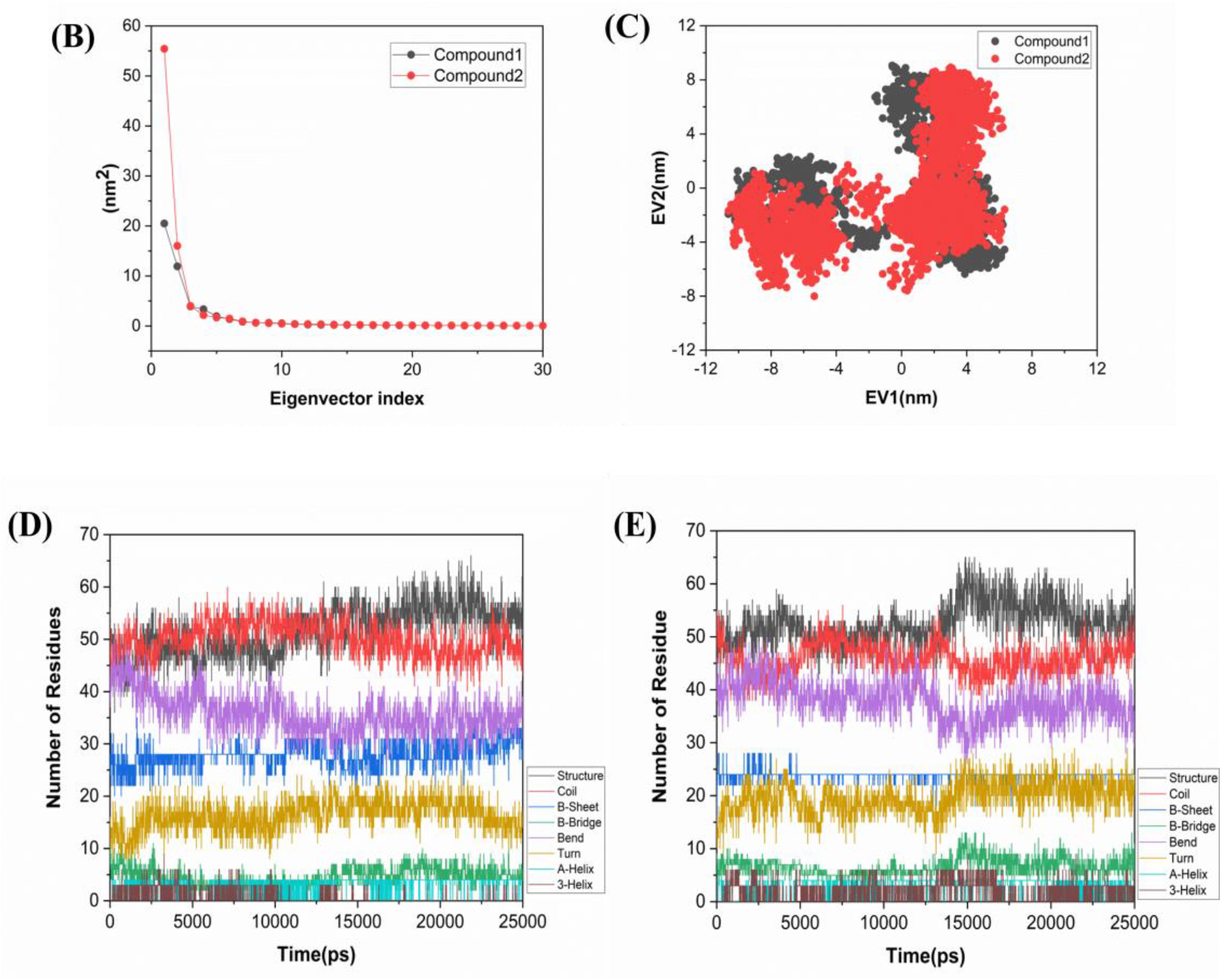
Hydrogen bond formation by rec-BSP1 with Compound 1(heparin, black) and Compound2 (PC, red) during 25 ns simulation. (A)Principal component analysis of rec-BSP1 with heparin and PC complexes. (B) The plot displaying the eigenvalue of both Compound 1 (heparin, black) and Compound2 (PC, red) of the first 30 eigenvectors. (C)The 2D Projection motion of the Compound 1 (heparin, black) and Compound2 (PC, red) in phase space along the first two principal eigenvectors. Secondary structure changes during the course of 25 ns MD simulation in (D) Heparin; (E) PC.

### Binding Free Energy Calculation

The binding free energy of rec-BSP1-ligand complexes were calculated using g_mmpbsa tool by molecular mechanics-based Poisson–Boltzmann Surface Area (35). In this study, 250 snapshots were taken from 25 ns MD trajectories and estimated the binding free energy for both complexes. The binding energy of rec-BSP1-heparin and rec-BSP1-PC complexes is −126.937 ± 95.414 KJ/mol, −45.485 ± 29.332 KJ/mol. The vdW energy was calculated with higher negative value of rec-BSP1-heparin complex because of solvent accessible surface area energy (ΔGsasa) and highest free energy due to Vander Waal, electrostatic interaction signifying the considerable hydrophobic interaction and stability of the rec-BSP1-heparin complex compared to rec-BSP1-PC complex (Table.1).

**Table 1.** Binding free energies of Heparin and PC complexes.

| Energy components (kJ/mol) | Heparin | PC |
| --- | --- | --- |
| Van Der Waal energy ($\Delta G_{\text{vdw}}$) | $-151.065 \pm 103.024$ | $-82.019 \pm 22.942$ |
| Electrostatic energy ($\Delta G_{\text{ele}}$) | $-51.488 \pm 44.724$ | $-8.879 \pm 16.056$ |
| Polar solvation energy ($\Delta G_{\text{ps}}$) | $90.919 \pm 67.678$ | $55.579 \pm 50.188$ |
| SASA energy ($\Delta G_{\text{sasa}}$) | $-15.305 \pm 10.014$ | $-10.166 \pm 2.529$ |
| Binding energy ($\Delta G_{\text{binding}}$) | $-126.937 \pm 95.414$ | $-45.485 \pm 29.332$ |

### Protein expression and purification

The optimized sequence was amplified by designed primers set with standardized annealing temperature of 52^0^ C and size of 468 bp was observed on 1% agarose gel (Fig. 5A). The gene was cloned into pET28a vector containing sequences for T7 promotor/lac operator, a His-tag and NdeI/XhoI restriction sites. Further it was confirmed by RE digestion (Fig. 5 B), colonies PCR (Supplementary fig. S4) and sequencing (Fig. 5 C). The recombinant plasmids were transformed into E. coli BL21 (DE3) strain and induce for expression of rec-BSP1 with 1 mM IPTG at 16^0^C for 22 h. The expressed His-tagged BSP1 was purified by Nickel Ion affinity chromatography and confirmed on SDS PAGE as a protein band of ~17 kDa (Fig. 6 A, B, C). The specificity checks and quantification were done by Western blot and Bradford assay (Fig. 6 D, E).

**Figure5.**
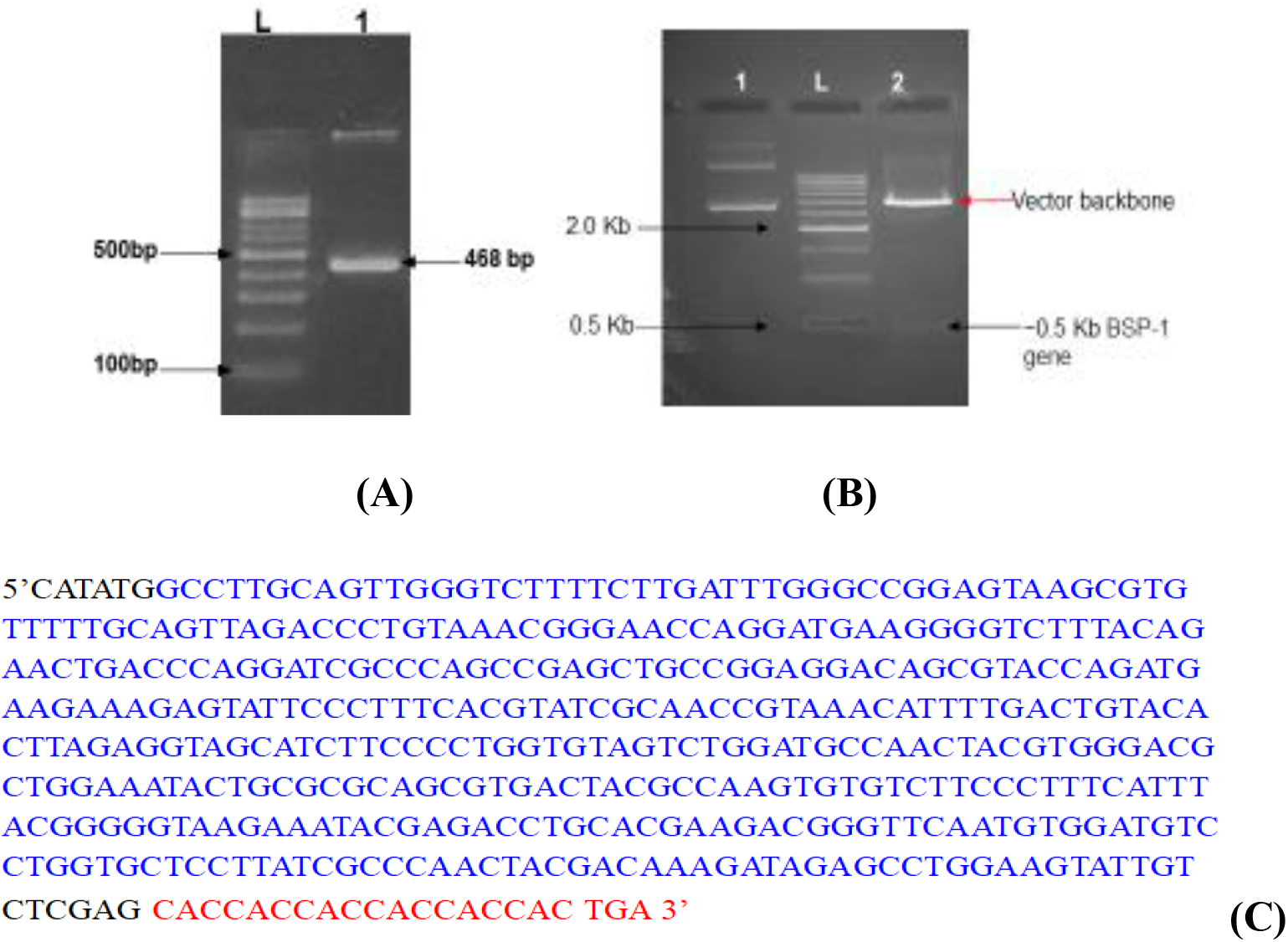
**(A)** PCR amplicon (lane1) loaded on 2% agarose gel, L: DNA ladder agarose gel. (B) Restriction enzyme NdeI/ XhoI loaded on 1.5% (C) Sequence of expression construct (rec-buBSP1) intact ORF with His Tag-Black Highlight-RE sites (NdeI/XhoI), Blue Highlight-rec-BSP1 gene, Red Highlight-His tag & stop codon.

**Figure6.**
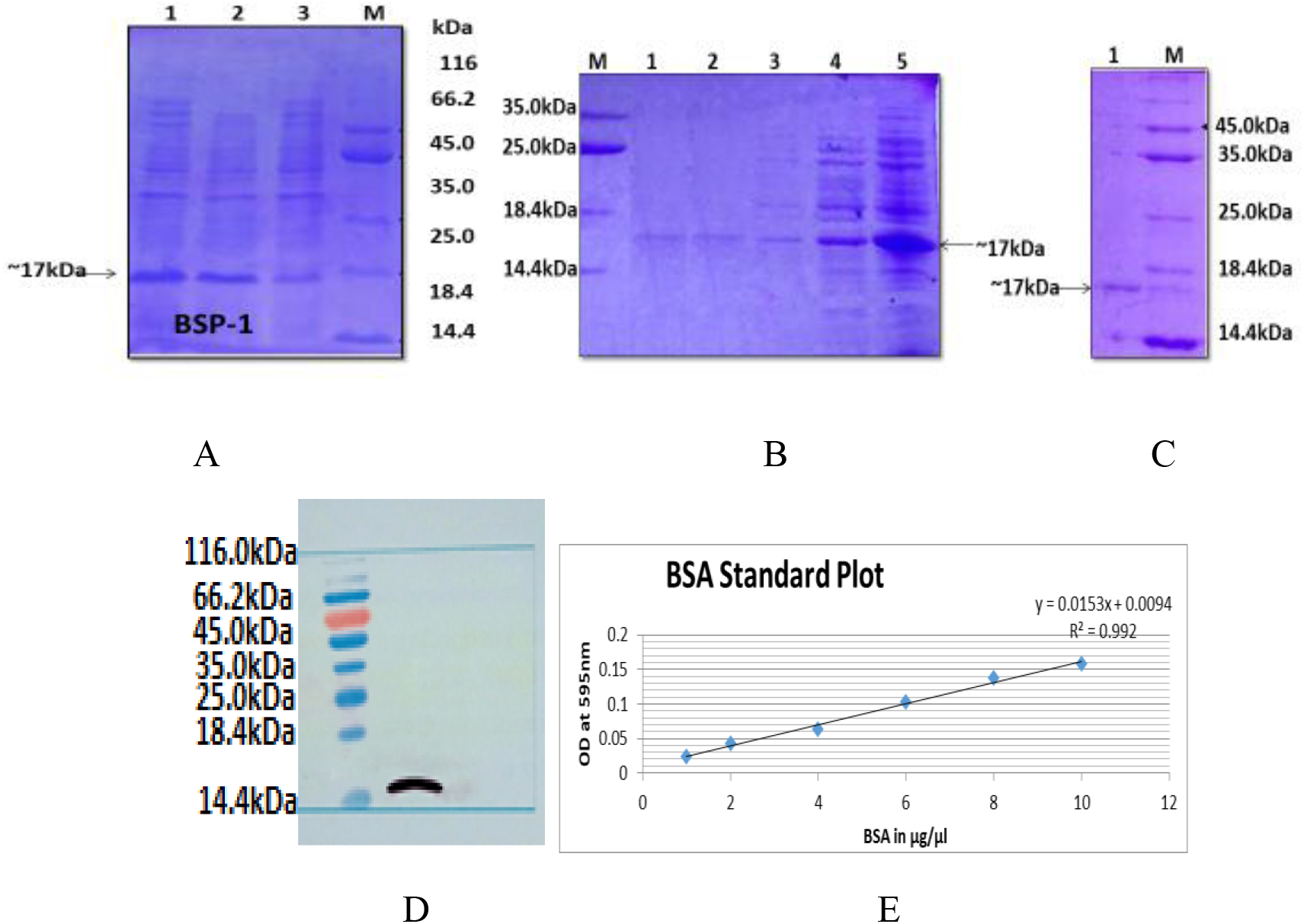
(A) Induction of PCR Positive Clones with 1mM IPTG at 16^0^C. Lane 1: 16 hours after induction sample (whole cell extract), Lane2: 4 hours after induction sample (whole cell extract), Lane3: Before induction sample (whole cell extract) showing slight leaky expression, Lane M: Protein Marker (B) Pellet and Supernatants after sonication were loaded on 15% SDS-PAGE where Lane 1: 1st sonication pellet, Lane 2: 2nd sonication pellet, Lane 3: 3rd sonication pellet, Lane 4: 4th sonication pellet, Lane 5: 4th sonication supernatant M: Marker (C) dialyzed pooled elutes loaded on 15% SDS-PAGE where Lane 1: Dialyzed pooled elutes; M: Marker. (D) Western Blot analysis of rec-BSP1 using anti his mouse Ab, HRP conjugate 2^0^ Ab showed ~17 kDa size protein for confirmation (E) Standard curve for estimation of rec-BSP1 concentration.

### Checking of motility percentage and acrosome reaction status of spermatozoa

The motility and acrosome reaction of spermatozoa were evaluated by computer assisted sperm analyzer (CASA), staining. It was observed a significant (P < 0.05) increase in percentage of motility and AR in sperm treated with 50 μg/ml of rec-BSP1 at 1 & 2hr and 2 &4 hr of incubation as compared to heparin/control (Fig. 7 A, B). The morphology of spermatozoa was found completely normal without any undesirable effect (Fig. 7 C, D).

**Figure7.**
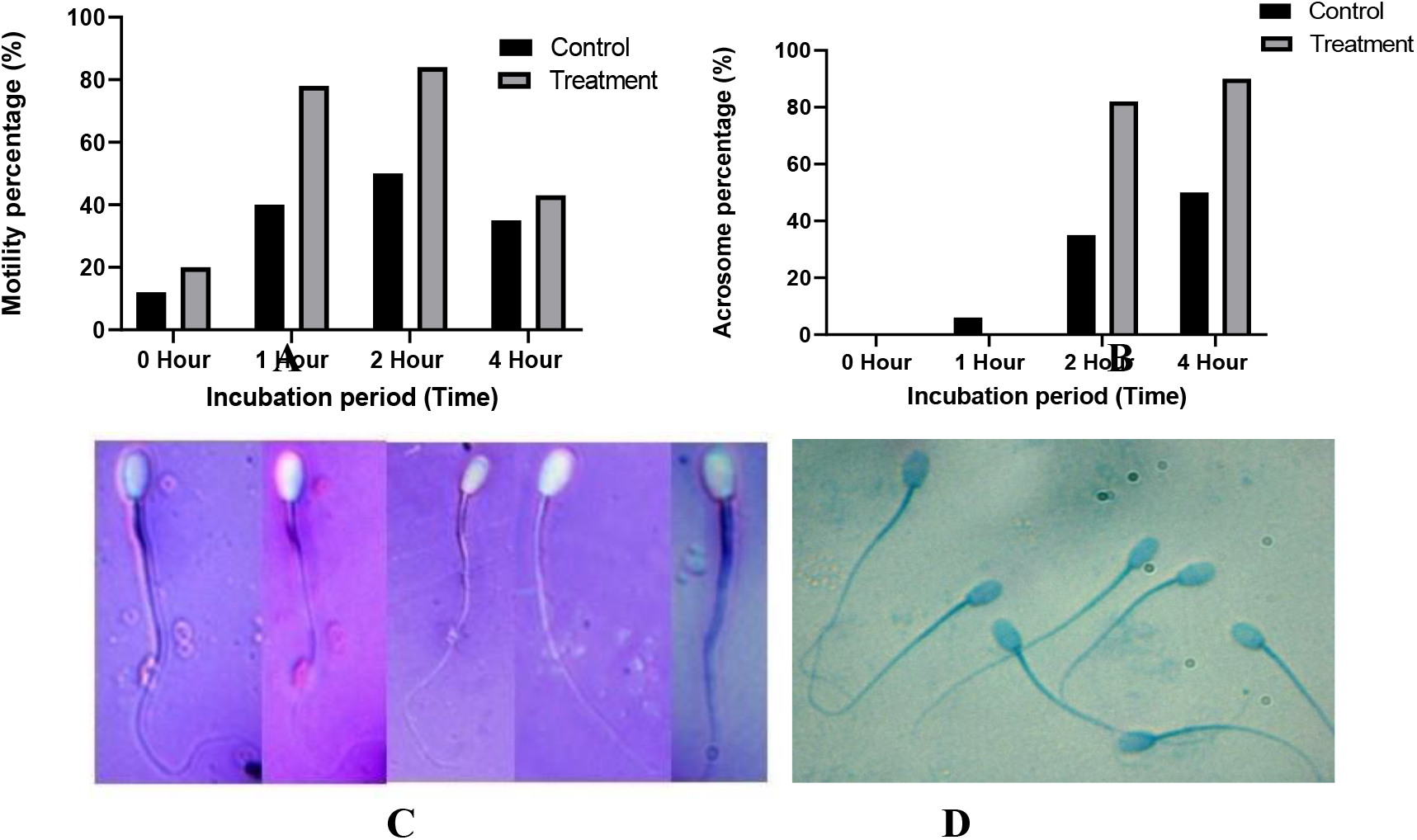
(A) Effect of 50 μg/ml rec-BSP1 and heparin in different incubation period (hours) on motility percentage of buffalos spermatozoa. (B) Effect of 50 μg/ml rec-BSP1 and heparin in different incubation period (hours) on acrosome percentage of buffalos spermatozoa. (C) Eosin Nigrosin staining of buffalo spermatozoa; all live sperm head clearly was clearly visible and showing bright head in 50 μg rec-BSP1 treatment group. (D) Trypan blue staining of buffalo spermatozoa; all sperm head was clearly visible and showing bright head in 50 μg rec-BSP1 treatment group.

### Effect of rec-BSP1 on *in vitro* embryo production

The effect of rec-BSP1 on *in vitro* buffalo embryo production was carried out with treatment of 20, 50 and 100 μg/ml of rec-BSP1 and heparin at concentration of 20 μg/ml and different stages of embryos (Fig.8). It was observed that the cleavage, blastocyst and hatched blastocyst production rate were significantly higher percentage in 50 μg/ml compare to 20 and 100 μg/ml of rec-BSP1 and control (heparin) (36) (Table.2). Further, hatched blastocysts were taken from 50 μg/ml rec-BSP1 treated group and control group for total cells counting by Hoechst 33342 staining. It was observed that there was significant increase of total cells number in hatched blastocysts in 50 μg/ml rec-BSP1 treated group compared to control (Fig. 9).

**Fig: 8.**
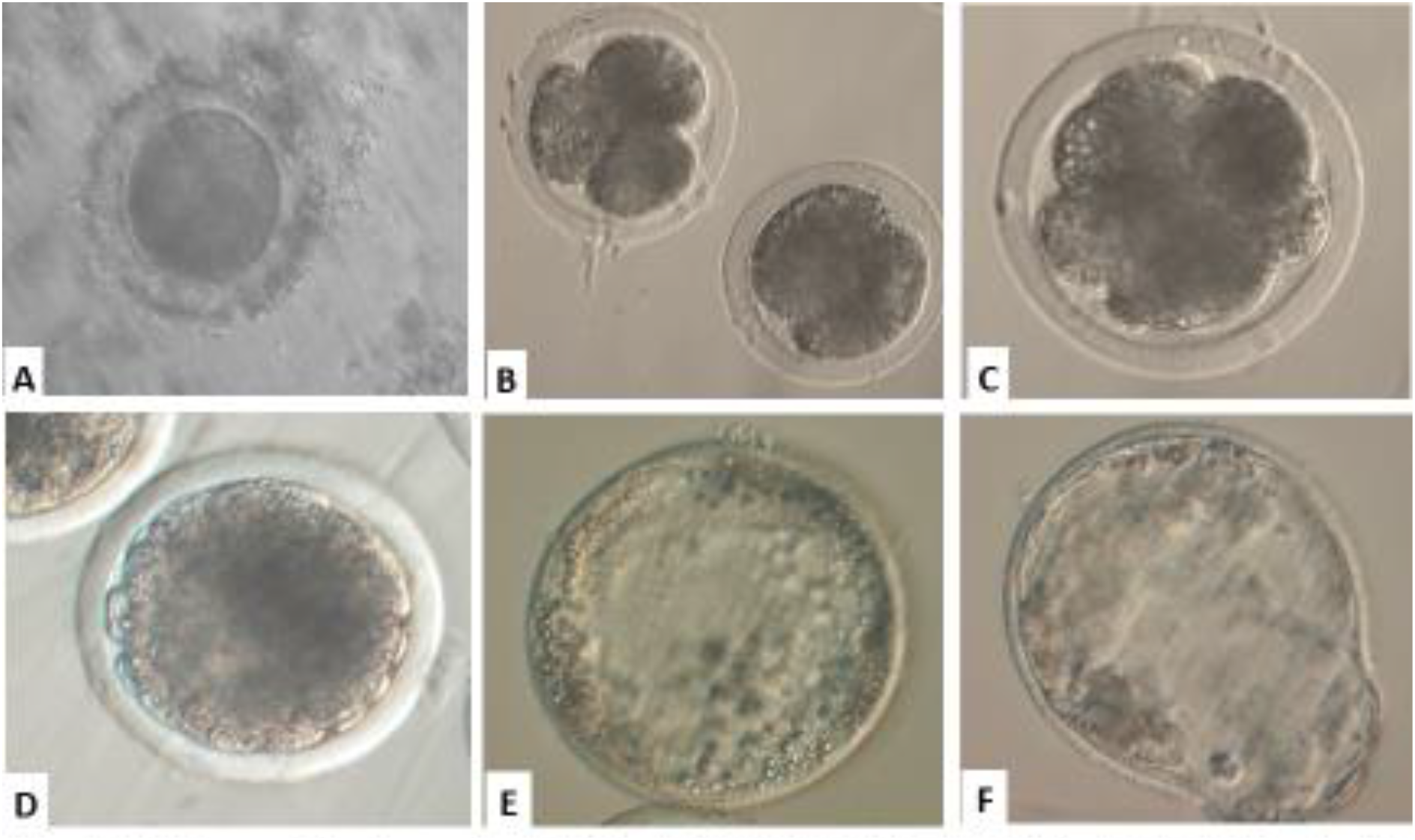
(A) Sperm-Oocyte co-incubation, (B) 2-Cell & 4-Cell, (C) 8-cell, (D) Morula, (E) Early Blastocyst, (F) Hatching blastocyst

**Figure 9.**
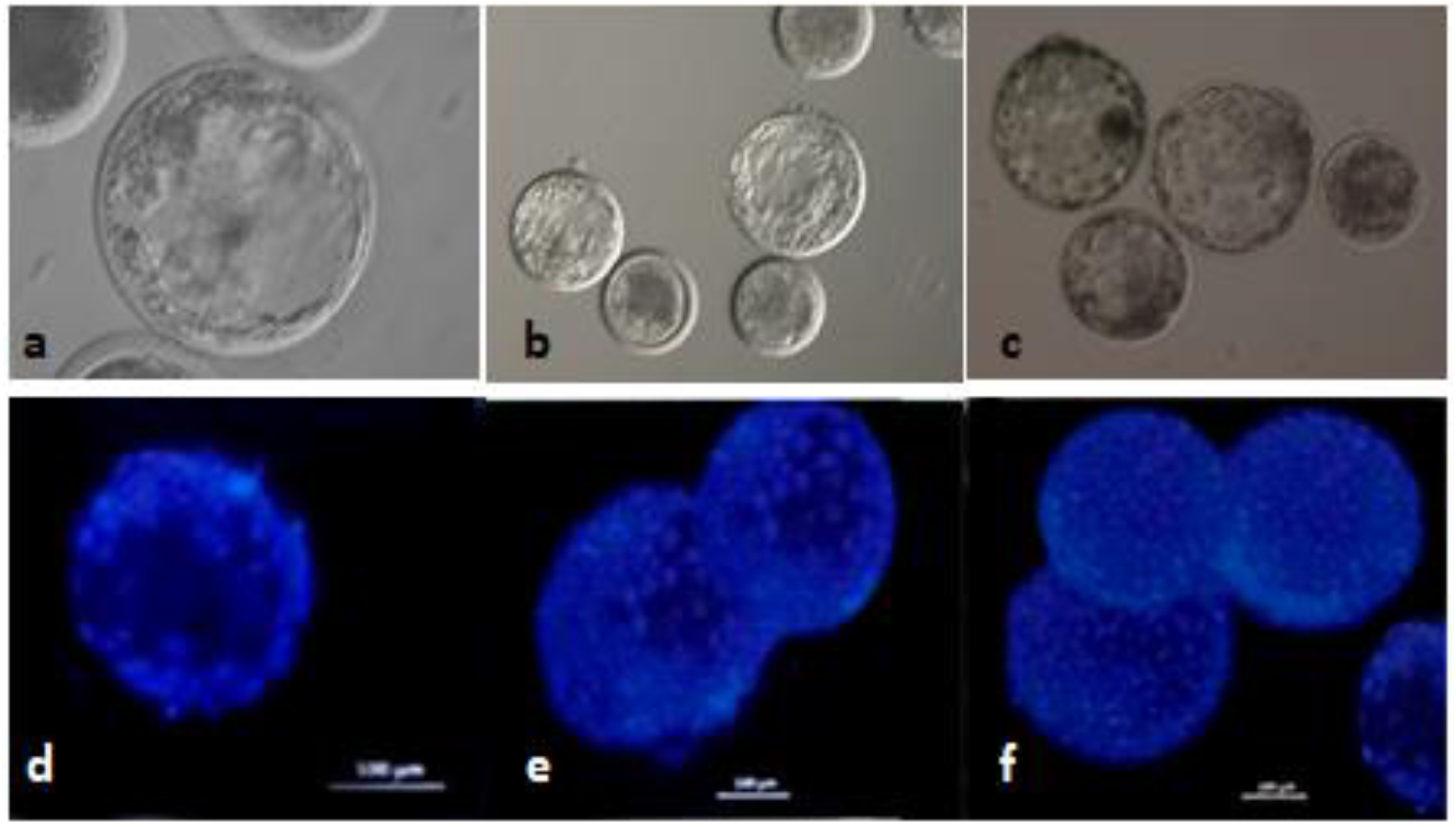
A few representative images of blastocyst and Hoechst staining of hatched blastocyst. Hatched /expanded /early blastocyst in control (a). Hatched blastocyst in IVF medium supplemented with 20 μg/ml of rBSP1 (b). Hatched blastocyst in IVF medium supplemented with 50 μg/ml of rBSP1 (c). Hoechst staining of blastocyst in control (d). Hoechst staining of blastocyst IVF (BO) medium supplemented with 20 μg/ml of rec-BSP1 (e). Hoechst staining of blastocyst IVF (BO) medium supplemented with 50 μg/ml of rec-BSP1 (f).

**Table 2.** Effect of rec-BSP1 on *in Vitro* Embryo production

| Concentration<br>rec-BSP1 µg/ml | No. of<br>oocytes | Cleavage rate<br>(%) | Blastocyst rate<br>(%) | Hatched Blastocyst<br>(%) |
| --- | --- | --- | --- | --- |
| Control (Heparin) | 303 | 177 (58.41) | 34 (19.21) | 31 (16.95) |
| 20 | 294 | 160 (54.42) | 24 (15.0) | 20 (12.50) |
| 50 | 305 | 198 (64.91) | 51 (25.75) | 44 (22.22) |
| 100 | 300 | 140 (46.66) | 16(11.43) | 13(9.28) |

## DISCUSSION

The mRNA sequence of *BSP* (*<u>Bubalus bubalis</u>*) was taken from the NCBI database. The sequences were aligned by the Muscle program using the UPGMA method and MegaX software was used to reconstruct the phylogenetic tree by the maximum likelihood method. It was observed in the phylogenetic tree that the recombinant protein having 99% similarity with bovine seminal plasma protein PDC 109 (OS: *Bos Taurus*). Three Dimensional Modeling of *rec-BSP1 was performed* and further structure Validated of *rec-BSP1* protein sequence and observed 138 amino acids. The active binding site of *rec-BSP1* was not known and probable binding sites of *rec-BSP1* were predicted by the software Metapocket. Molecular docking analysis of *rec-BSP1* was done using two ligands of heparin and phosphatidylcholine using AutoDock 4.2.6 tool. Molecular Docking simulation study was also carried out using GROMACS v 5.0.7 package under Amber99B-ILDN force field for understanding the dynamics and structural behavior of the *rBSP1* with the two ligands of heparin and phosphatidylcholine. The hydrogen bond was analyzed between *rBSP1* and the ligands and calculated for the overall stability of protein-ligand complexes using gmx bond over 25 ns simulation period. The sequence was optimized and designed the primers with a standardized annealing temperature of 52^0^ C and a size of 468 bp was observed and cloned the gene into cloning vectors. Then the cloned gene was further cloned into a pET28a expression vector containing sequences for T7 promoter/lac operator, a His-tag, and NdeI/XhoI restriction sites.

The rec-BSP1 expressed at 16^0^C was purified by Ni NTA affinity chromatography because 6 His tag fused with N terminal protein. The recombinant protein was bound and eluted from Ni NTA agarose bead using PH 4.5 buffer. The purified protein was visualized on SDS PAGE and further confirmed by Western blot analysis in which the membrane was incubated with anti-6x-His epitope tag mice (Thermofisher) at 1:1000 dilution. The secondary antibody HRPO conjugated Anti-Mouse IgG at 1:1000 dilutions and immunosignal was detected by 3, 3’-Diaminobenzidine (DAB) staining. We tried to examine its functional activities such as motility and morphology of spermatozoa by CASA study and staining (Eosin, nigrosine/trypan blue). The experiment revealed a significant increase in percentage motility and a significant number of spermatozoa with better morphology in the 50 μg/ml rec-BSP1 treated group as compared to the control. We developed a method for the expression of a soluble form of rec-BSP1 in the prokaryotic system and purified the rec-BSP1 which was confirmed by western blot. The rec-BSP1 was found to be effective for enhancing sperm motility and improving the morphology of spermatozoa and a significant increase in blastocyst production. Molecular docking and MD simulation experiments proofed that rec-BSP1 interacts with heparin with stronger affinity than PC and rec-BSP1 heparin complex is more stable than rec-BSP1 phosphatidylcholine complex. ProSA confirmed the consistency and robustness of the proposed homology model and heparin with the highest docking score seems to enhance the activity of the rec-BSP1 and crystal structure of seminal plasma protein PDC-109 as the best PDB template and need to be further analyzed.

## CONCLUSIONS

In conclusion, this study of the role of rec-BSP1 in reproductive function may recognize a new fertility factor that can be beneficial to enhance male fertility. Applicably, the research work suggests that the proposed homology model of heparin may a suitable, consistent, and robust model with the highest docking score seems to improve the activity of rec-BSP1 and ultimately use as a potent fertility factor which enhances *in vitro* embryo production.

## Acknowledgments

This work is part of PhD thesis of S. Bag, in Animal Biotechnology of ICAR-NDRI, Karnal (Deemed University), Haryana, India. The authors also acknowledge the Director, CCS National Institute of Animal Health, Baghpat, UP, Department of Animal Husbandry and Dairying, Government of India for providing necessary facilities for completion of the research work.

## Authors’ contributions

SB (Sudam Bag) performed most of the experiments, analysis data and manuscript preparation. SA, SS, production of buffalo blastocysts, AT, NCM, SB, SS, SKS, assist in experiments of cloning, expression, purification, and embryo production research work and data analysis, and PM, SK, DM have been involved in final data analysis, manuscript preparation, and presentation.

## Competing interests

All the authors declare that they have no conflict of interest.

## Ethical approval and informed consent

Ethical approval was taken from Institute ethics committee, ICAR-National Dairy Research Institute (NDRI), Karnal, India during the study. All experiments were conducted as per the rules, guidelines, and regulations of ICAR-National Dairy Research Institute (NDRI), Karnal, India Institute Animal Ethics Committee. This research work was approved by the Institute of Animal Ethics Committee, NDRI, India and all methods were performed in accordance with the relevant guidelines and regulations though no animal participant involved in this research.

---

**Supplementary S1.**
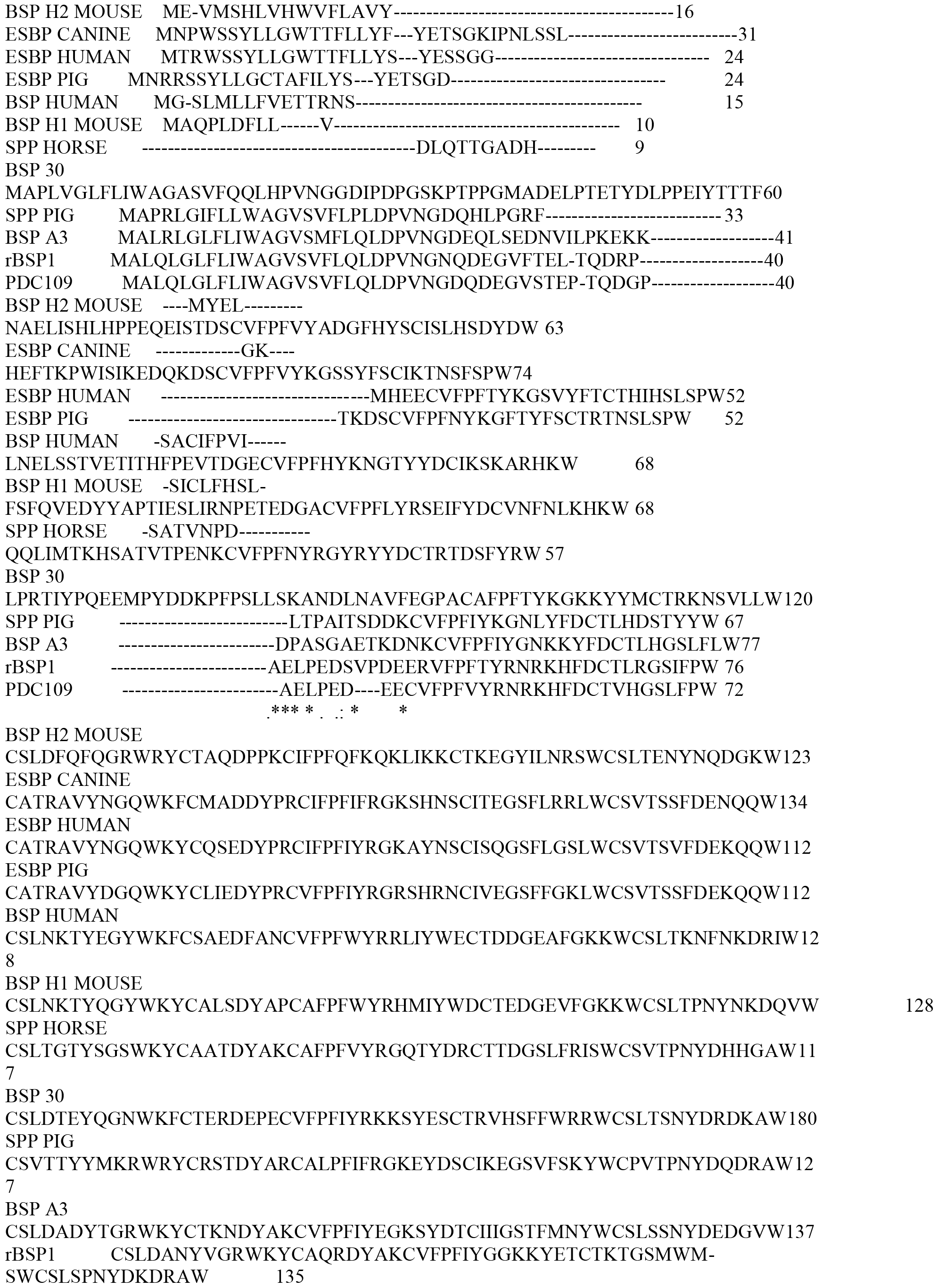
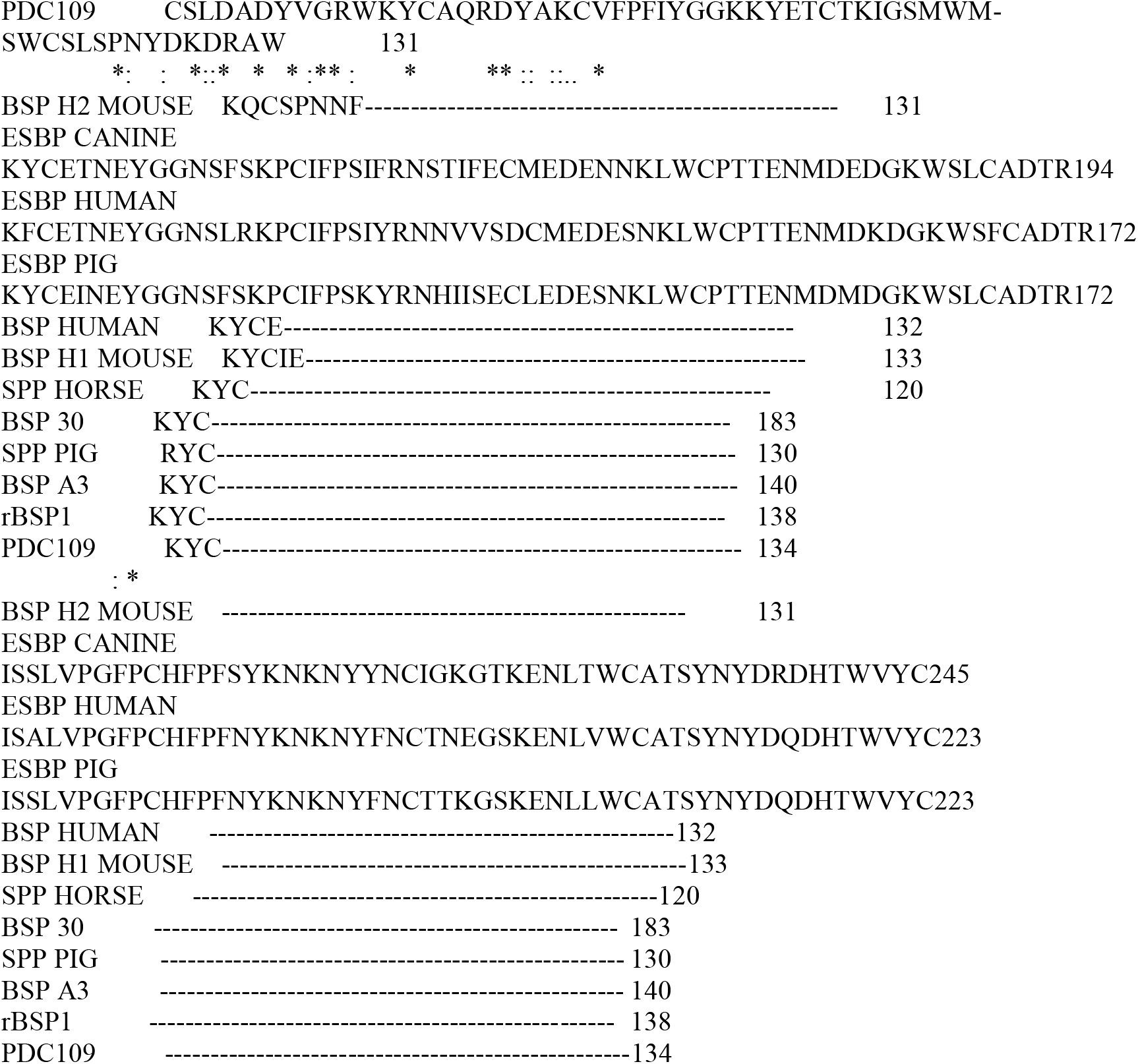

**Supplementary S2.**
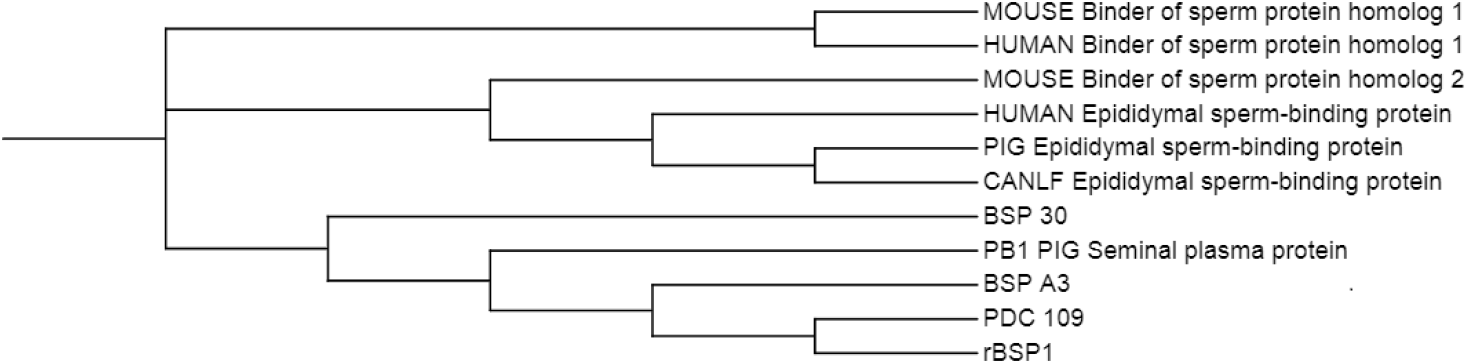
Multiple Alignments of Amino Acids Sequence of seminal vesicular protein of different species retrieved from GenBank. Residues Identical to the Sequences of Indicated by Dots (.). Spaces or Dash (-) Denotes Gap & star (*) Denotes conserved amino acid residues. Schematic Maximum Likelihood Consensus Tree with 1000 Boot straps. The Bootstrap Confidence value at the Nodes Represent the percentage of Times the Group occurred out of 1000 Trees & the Bar represents the Genetic Distance.

**Supplementary S3.** Binding energy, Ligand efficiency and Hydrogen bond formation of rBSP1 protein.

| SN | Compounds | Binding Energy<br>(kcal/mol) | Ligand<br>Efficiency | Hydrogen bond |  |
| --- | --- | --- | --- | --- | --- |
|  |  |  |  | Residues | Bonds |
| 1 | Heparin | -5.44 | -0.25 | ASN 26<br>THR 33<br>ARG 53<br>ASP 66 | 4 |
| 2 | Phosphatidyl<br>Choline | -5.21 | -0.07 | THR 33<br>GLU 34 | 2 |

**Supplementary S4.**
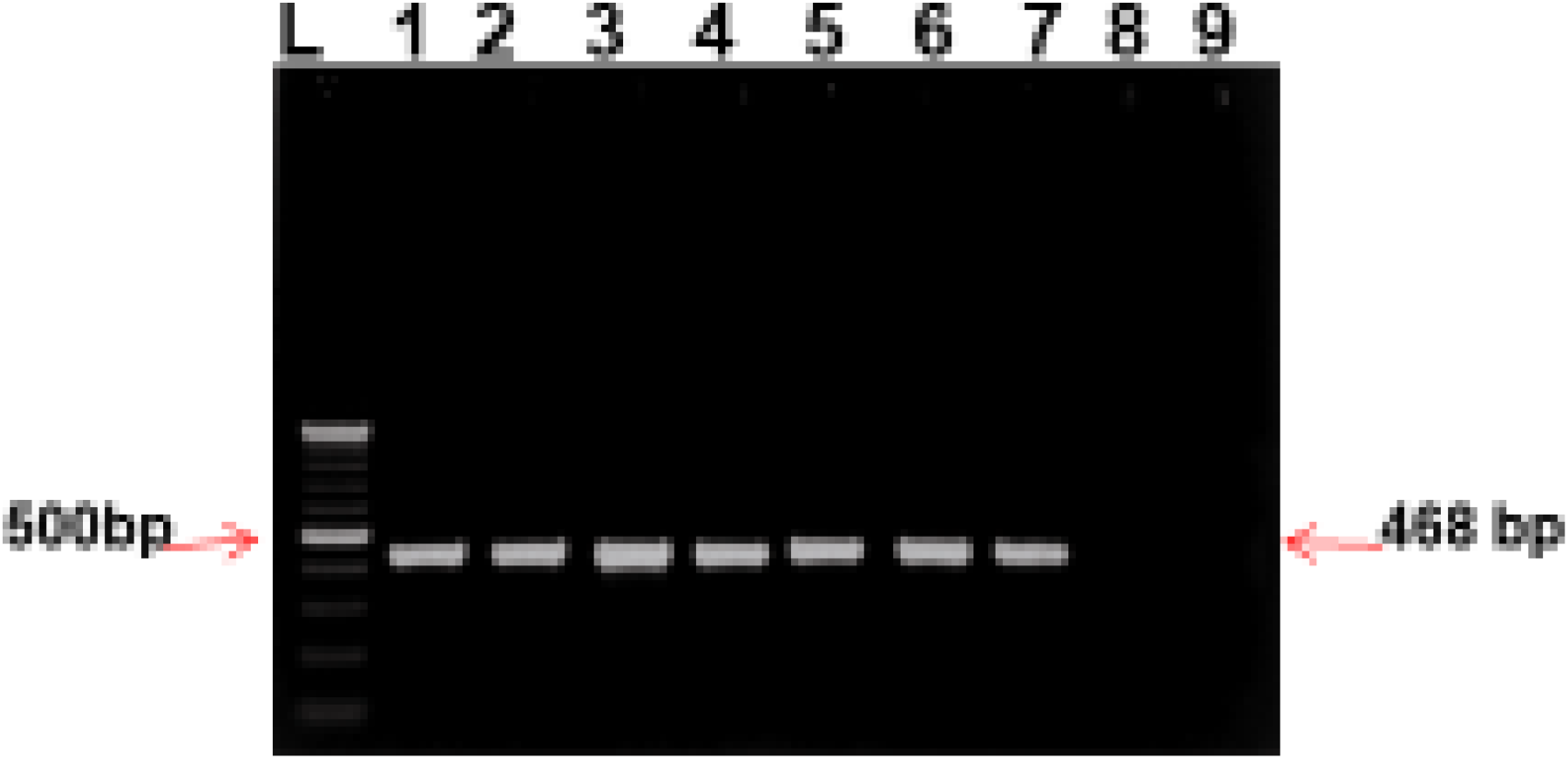
Colony PCR of BSP gene where Lane M: DNA Ladder; Lane 1-7: Positive clones.

## Notes

### Competing Interest Statement

The authors have declared no competing interest.

